# Narrow equilibrium window for complex coacervation of tau and RNA under cellular conditions

**DOI:** 10.1101/424358

**Authors:** Yanxian Lin, James McCarty, Jennifer N. Rauch, Kris T. Delaney, Kenneth S. Kosik, Glenn H. Fredrickson, Joan-Emma Shea, Songi Han

**Affiliations:** Biomolecular Science and Engineering, University of California Santa Barbara, Santa Barbara, CA 93106, USA; Department of Chemistry and Biochemistry, University of California Santa Barbara, Santa Barbara, CA, 93106, USA; Molecular, Cellular and Developmental Biology, University of California Santa Barbara, Santa Barbara, CA 93106, USA; Neuroscience Research Institute, University of California Santa Barbara, Santa Barbara, CA 93106, USA; Materials Research Laboratory, University of California Santa Barbara, Santa Barbara, CA 93106, USA; Department of Chemical Engineering, University of California Santa Barbara, Santa Barbara, CA 93106, USA; Department of Physics, University of California Santa Barbara, Santa Barbara, CA 93106, USA

## Abstract

The conditions that lead to the liquid-liquid phase separation (LLPS) of the tau protein, a microtubule associated protein whose pathological aggregation has been implicated in neurodegenerative disorders, are not well understood. Establishing a phase diagram that delineates the boundaries of phase co-existence is key to understanding its LLPS. Using a combination of EPR, turbidity measurements, and microscopy, we show that tau and RNA form complex coacervates with lower critical solution temperature (LCST) behavior. The coacervates are reversible, and the biopolymers can be driven to the supernatant phase or coacervate phase by varying the experimental conditions (temperature, salt concentration, tau:RNA charge ratio, total polymer concentration and osmotic stress). Furthermore, the coacervates can be driven to a fibrillar state through the addition of heparin. The equilibrium phase diagram of the tau/RNA complex coacervate system can be described by a Flory-Huggins model, augmented by an approximate Voorn Overbeek electrostatic term (FH-VO), after fitting the experimental data to an empirical Flory interaction parameter divided into an entropic and enthalpic term. However, a more advanced model in which tau and RNA are treated as discrete bead-spring chains with a temperature-dependent excluded volume interaction and electrostatic interactions between charged residues, investigated through field theoretic simulations (FTS), provided direct and unique insight into the thermodynamic driving forces of tau/RNA complexation. FTS corroborated the experimental finding that the complex coacervation of tau and RNA is has an entropy-driven contribution, with a transition temperature around the physiological temperature of 37 °C and salt concentrations around 100-150 mM. Together, experiment and simulation show that LLPS of tau can occur under physiological cellular conditions, but has a narrow equilibrium window over experimentally tunable parameters including temperature, salt and tau concentrations. Guided by our phase diagram, we show that tau can be driven towards LLPS under *live* cell coculturing conditions with rationally chosen experimental parameters.

## Introduction

Protein liquid-liquid phase separation (LLPS) is a process in which proteins assemble and partition into a protein-dense phase and a protein-dilute phase. The proteins in the dense phase form droplets, and retain liquid-like mobility, as shown by NMR measurements [1,2]. The process of LLPS *in vitro* has been observed for decades [3–8], but the field has recently been invigorated by the realization that LLPS also occurs *in vivo*, suggesting a possible physiological role for these assemblies[4,9,10]. The overwhelming majority of proteins observed to undergo LLPS are intrinsically disordered proteins (IDPs) [11], and much of the research thus far has focused on ALS-related IDPs, including FUS [9,12–14], hnRNPA2B1 and hnRNPA1 [15], TDP-43 [15,16], C9ORF72 [17–19] and Ddx4 [20]. Recently, we and others discovered that another amyloid forming IDP, the microtubule binding protein tau, also undergoes LLPS [21–25]. Interestingly, many of the LLPS forming IDPs have been observed to form amyloid fibrils in cell-free systems [13,15], leading to a number of hypotheses regarding the physiological role of LLPS in regulating aggregation. In particular, a compelling idea is that protein LLPS may be an intermediate regulatory state, which could redissolve into a soluble state or transition to irreversible aggregation/amyloid fibrils [13–15,21,22].

In a healthy neuron, tau is bound to microtubules. When tau falls off the microtubule under adverse conditions to the cell, tau is solubilized in the intracellular space as an IDP. Under certain conditions, tau forms intracellular fibrillary tangles, a process linked to neurodegenerative tauopathies that include Alzheimer’s disease. In recent work, we showed that tau in neurons strongly (nanomolar dissociation constant) and selectively associates with smaller RNA species, most notably tRNA [22]. We also found tau and RNA, under charge matching conditions, to undergo LLPS [22] in a process determined to be complex coacervation (CC) [26]. We found that tau-RNA LLPS is reversible, and persisted for > 15 hours without subsequent fibrilization of tau, and hypothesized that LLPS is potentially a physiological and regulatory state of tau.

In this work, we characterize the phase diagram of tau/RNA LLPS using a combination of experiment and simulation, and thereby specify the conditions that drive the system towards a homogeneous phase or an LLPS state. We study a N-terminus truncated version of the longest isoform of human 4R tau *in vitro*, and first demonstrate that tau/RNA complexation is reversible, and that tau remains dynamic and without a persistent structure within the dense phase. The phase coexistence curve separating a supernatant phase from a condensate phase is determined by the system’s free energy, which in turn is state dependent, *i.e*. dependent on concentration, temperature, salt, and the nature of the interaction strength between the various solution constituents, including the solvent. We construct the phase diagram from cloud-point measurements of the onset of complex coacervation under varying conditions of temperature, salt, and polymer concentrations. These experiments establish the features and phase coexistence boundaries of the phase diagram, which we then model using theory and simulation to rationalize and understand the physical mechanisms that drive and stabilize LLPS.

A number of theoretical models can be used to model LLPS, each with their own advantages and disadvantages. Ideally, one would turn to simulations at atomic resolution in explicit solvent; however, such models are computationally prohibitive given the multiple orders of magnitude in time and length scales involved in LLPS. Turning to the polymer physics literature, theoretical treatments of simplified coarse-grained models are much more computationally tractable, and offer useful insight. Although approximate, analytical theories can be formulated, providing an extremely efficient platform for describing the thermodynamics of polyelectrolyte mixtures [27]. These include the Flory-Huggins model [28], the Voorn-Overbeek model [6,29–34], the random-phase approximation [35–37], the Poisson-Boltzmann cell model [38,39], as well as other more sophisticated approaches[40–42], which have been applied to synthetic polymers with low sequence heterogeneity [29,43–46], and to proteins with single composition [2,20,47,48]. While such models have been successful in describing simpler polyelectrolytes, it is less apparent that these models are suitable to describe the complex coacervation of the more complicated tau/RNA system. The simplest approach that one can use is the Flory-Huggins (FH) model, augmented by the Voorn and Overbeek (VO) correction to describe electrostatic correlations. This model is widely used to model LLPS; however, while experimental data can be fit to the model [2,20], ultimately the FH-VO model has serious inadequacies. The original Flory-Huggins model is a mean-field theory, which means that fluctuations in polymer densities away from their average value in each phase are neglected. Augmenting the FH model with a VO treatment of electrostatics approximately accounts for charge correlations, but it entirely neglects chain-connectivity [49]. Thus, the FH-VO model is unable to model the spatially varying charge distribution along the polymer backbone. Ideally, one would like to introduce chain connectivity, charge correlation, and uneven charge distribution into a more realistic polymer physics model; however, a full treatment of polymer density fluctuations is analytically intractable. One possible approach is to pursue a Gaussian approximation to field fluctuations, also known as the random phase approximation (RPA) [50–52]. The RPA model can be viewed as a lowest-order correction to the mean field approximation, and was recently introduced to describe the charge pattern and sequence-dependent LLPS of IDPs [53,54]. The advantage of the RPA model, over the mean-field FH-VO model, is that charge correlations are introduced in a formally consistent manner. Nonetheless, it has been recently demonstrated that the RPA model fails to quantitatively predict polymer concentrations in the dilute phase, given that higher-order fluctuations are important in this regime [55,56].

Of all the models described above, fitting experimental data with the FH or FH-VO theory is currently the preferred methodology in the LLPS community to describe and analyze phase diagrams. We demonstrate that this model can be fit to describe our experimental data, but the learning outcome from this modeling is limited. Thus, we take a different approach by computing the exact phase diagram of an off-lattice coarse-grained polyelectrolyte model using field theoretic simulations (FTS). FTS is a numerical approach that allows one to fully account for fluctuations, and thus serves as an approximation-free method to compute equilibrium properties from a suitably chosen coarse-grained representation of the true system. The ability to perform field theoretic simulations enables us to include the important physics of polymer sequence-specificity that cannot be captured by FH-VO, including charge distribution and chain connectivity. Results from FTS are compared to those obtained from the FH-VO model.

The model substantiates the experimental phase diagram that the equilibrium window for the complex coacervation of tau and RNA under cellular conditions is narrow. Guided by the phase diagram, empirically obtained from *in vitro* experiments and validated by simulation, we finally show that LLPS of tau-RNA can be established and rationalized under cellular co-culturing conditions in the presence of live cells.

## Results

### Tau-RNA complex coacervate is reversible and a dynamic liquid phase

Truncated versions of the longest isoform of human 4R tau, residues 255-441 [57] and residues 255-368 were used to study tau-RNA complex coacervation (CC). A C291S mutation was introduced to either tau variant, resulting in single-cysteine constructs. Thioflavin T assays and TEM imaging were performed showing these variants retain the capability to form fibrils with morphology similar to full length tau. Unless otherwise specified, we refer to these two single-cysteine tau constructs as tau187 and tau114 (tau114 is close to K18, 244-372 [58]), respectively, while tau refers collectively to any of these variants (see SI methods for experimental details). Importantly, experiments were performed with freshly eluted tau within 30 minutes upon purification to minimize the effects of possible disulfide bond formation. This minimizes the influence of the cysteine mutations on the LLPS behavior of tau-RNA CC. The single-cysteine containing tau187 can be singly spin labeled at site 322, referred to as tau187-SL (see SI methods). Full length tau, tau187 and tau114 are overall positively charged with an estimated +3, +11 and +11 charge per molecule at neutral pH, respectively, based on their primary sequences. The charged residues of tau are more concentrated in the four repeat domains (Fig 1A). PolyU RNA (800~1000 kDa), which is a polyanion carrying 1 negative charge per uracil nucleotide, was used in this study and henceforth referred to as RNA (Fig 1A). Under ambient conditions, both tau and RNA are soluble and stable in solution. By mixing tau and RNA under certain conditions, a turbid and milky suspension was obtained within seconds, where tau and RNA formed polymer-rich droplets (dense phase) separated from polymer-depleted supernatants (dilute phase) (Fig 1B). These polymer-rich droplets are tau-RNA CCs. We began by determining the concentration of the dense and dilute phases. After mixing and centrifuging 60 µL tau187-RNA droplet suspension, we separated a polymer-rich phase of volume <1 µL with a clear boundary against the dilute supernatant phase. Applying UV-Vis spectroscopy (see SI methods), we determined the concentration of tau and RNA inside the droplets as >76 mg/mL and >17 mg/mL with partitioning factors of >15 and >700 respectively. This is consistent with our previously findings that tau is virtually exclusively partitioned within the dense phase [22]. High protein concentrations are typically correlated with higher propensity for irreversible protein aggregations. In order to verify that there was indeed no fibril formation, tau187-RNA CCs were prepared by mixing tau187-SL and RNA (see SI methods) and monitored by continuous wave electron paramagnetic resonance spectroscopy (For details of cw-EPR experiments see SI methods). The cw-EPR spectra shows no broadening (Fig 1C), and the cw-EPR spectra analysis reveals an unchanged rotational correlation time for the spin label of tau187-SL, τ, of 437 ± 37 ps as a function of time after > 96 hours of incubation at room temperature (Fig 1D, turquoise) (see SI methods). For comparison, cw-EPR spectra and τ were recorded of tau187-SL alone in buffer, and of tau187-SL in the presence of heparin under fibril forming conditions. Tau187-SL alone in buffer showed cw-EPR spectra overlapping with those of tau187-RNA CC, and rotational correlation time τ, 425 ± 16 ps, nearly identical to the τ of tau187-SL CCs (Fig. 1D, red). In contrast, tau187-SL with heparin shows a significantly broadened cw-EPR spectrum and an increasing τ to 2.3 ± 0.7 ns (Fig 1C, D, green). Note that a hundreds of ps range of τ corresponds to rapid tumbling of the spin label, whose rotational degree of freedom is minimally hindered by molecular associations, while a several ns range of τ corresponds to slow tumbling and molecular hindering by association or confinement. The Thioflavin T (ThT) fluorescence curves of the same sample system as a function of time confirms the absence of amyloid aggregate formation in tau187-RNA CCs (Fig S1). These results together suggest that tau187-RNA CCs are in an equilibrium state, in which tau retains its solution-like dynamics.

**FIG. 1.**
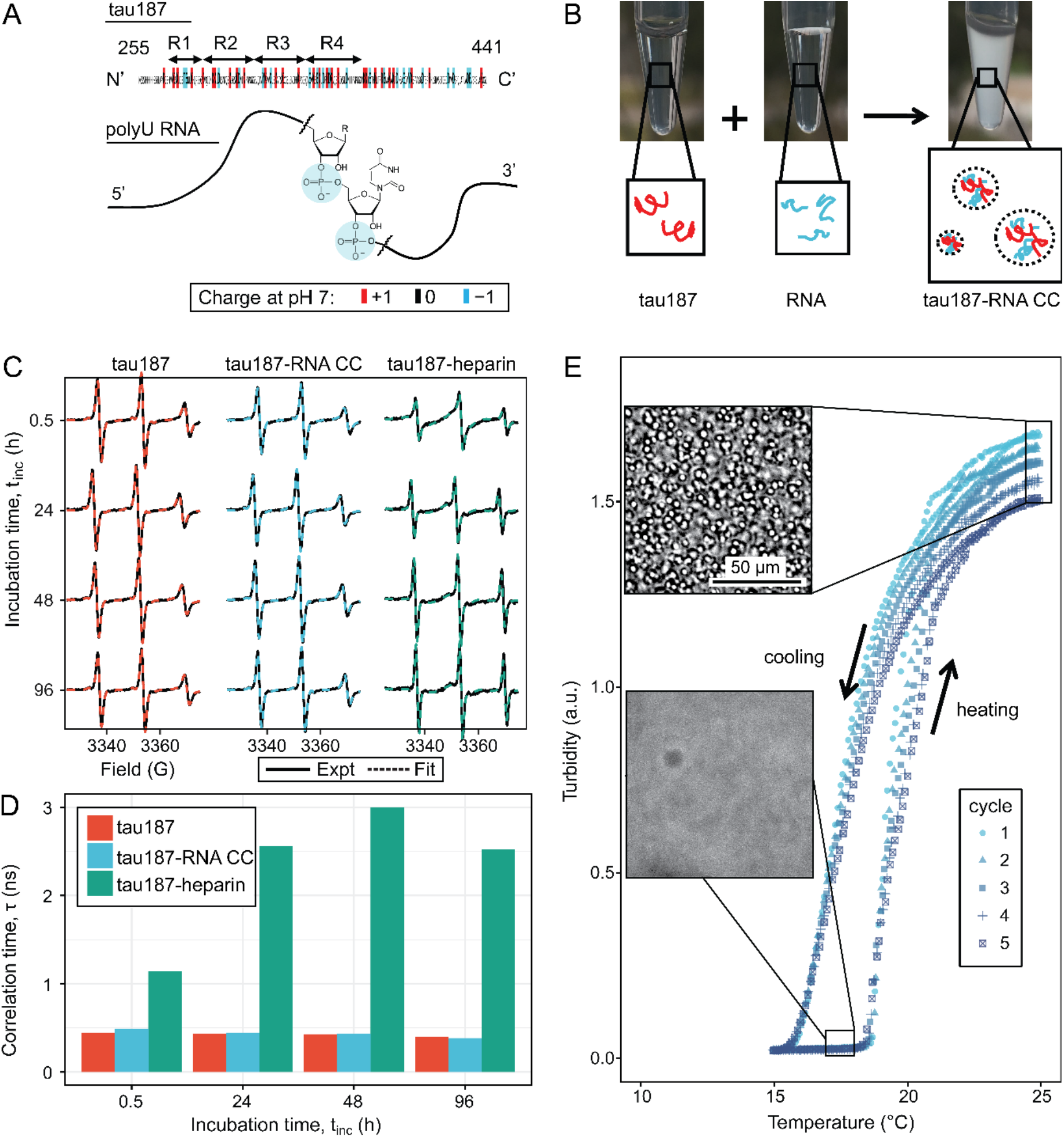
Steady tau dynamics and reversible droplet formation of tau-RNA complex coacervates. **A.** Diagram of tau187 (tau) and polyU RNA (RNA). Tau187 is a truncated version of full-length human tau (2N4R 255-441) containing repeat domains and C terminal. At neutral pH experimental conditions, tau187 is overall positively charged; while RNA consists of a phosphate backbone and is negatively charged. **B.** Scheme of tau-RNA CC preparation. Mixing clear tau187 and RNA solutions at proper conditions results in a turbid solution containing liquid droplets. **C.** X-band cw-EPR spectra (solid line) of tau187 solution (tau187, red), tau187-RNA CC (blue) and tau187-heparin (green) at room temperature with different incubation time, t_inc_. Samples contains 500 µM tau with 20% spin-labelled. EPR simulation were performed (SI Method) and the fitted spectra is shown as a dashed line. **D.** Rotational correlation time, τ_R_ extracted from EPR simulation shown in (b1) (SI Method). **E.** Turbidity of tau187-RNA suspension in consecutive heating-cooling cycles. Confocal images represented samples at 19 °C and 25 °C. Temperatures were ramped at 1 °C/min.

Next, we investigated the reversibility of tau187-RNA complex coacervation. Tau187-RNA CCs were prepared again and incubated by cyclically ramping the temperatures (1 °C/min) upwards and downwards, while the absorbance at λ = 500 nm was monitored, referred to as turbidity hereafter. Ramping rates of 0.5 °C/min and 1 °C/min were tested, but the results shown to be indistinguishable. Microscopy images were concurrently acquired at low and high turbidity, confirming the appearance and abundance of CC droplets correlating with turbidity increase, and *vice versa* (Fig 1E). The turbidity-temperature curves show that at high temperature, samples became turbid with Abs_500_ ~ 1.5 and abundance of CCs, while at low temperature, samples became transparent with Abs_500_ ~ 0 and absence of CCs. This demonstrates tau187-RNA CC formation is favored at higher temperature, following clearly a lower critical solution temperature behavior (LCST) (Fig 1E) [59]. By cycling the temperature, we robustly and reversibly changed the tau187-RNA mixture between a turbid state to a completely transparent state (Fig 1E). The transition temperatures at which the turbidity emerged during heating and vanished during cooling stay invariant with repeated heating-cooling cycles. The method of extracting a cloud point for the LCST transition temperature from such data will be described in detail in the next section. Importantly, the history of temperature change does not affect the resulting state. Hence the formation and dissolution of tau187-RNA CCs are reversible and consistent with a path-independent equilibrium process. We point out that the maximum turbidity value successively decreases with each heating cycle (Fig 1E), even though the transition temperatures remain invariant. This can be attributed to slow degradation of RNA with time, (as demonstrated in Fig S2) by verifying an altered turbidity change in the presence of RNase or RNase inhibitor.

It is understood that upon gradual heating of the solution phase, the mechanism of LLPS proceeds via a nucleation process [60], and hence there is a kinetic barrier evidenced by the observed hysteresis in Fig 1E. Nonetheless, we conclude that the final tau-RNA CC state reached upon heating is a true thermodynamic state, and thus can be modeled by an equilibrium theory of phase separation.

### Tau-RNA complex coacervate phase diagram

To understand the principles and governing interactions driving tau-RNA CC formation, we constructed a phase diagram for tau187-RNA CC by measuring the transition temperature – to be described in greater detail below – as a function of protein concentration and salt concentration. We first recorded tau187-RNA turbidity at various [tau], [RNA] and [NaCl] values, ranging from 2-240 µM, 6-720 µg/mL and 30-120 mM, respectively. Titrating RNA to tau187, the turbidity was found to be peaked when [RNA]:[tau] reached charge matching condition at which the charge ratio between net positive and negative charges was 1:1 (which for tau187 and RNA used in this study corresponded to [tau187]:[RNA] = 1 µM : 3 µg/mL), validating once more that LLPS is driven by complex coacervation (CC) (Fig S3). Henceforth, all phase diagram data are acquired at a charge matching condition between RNA and tau. Titrating NaCl to tau187-RNA, CC formation showed a steady decrease of turbidity (Fig S3). Combined, these demonstrate that tau187-RNA CC favors the condition of charge balance and low ionic strength, which is consistent with known properties of CC and previous findings [22].

We next investigated the phase separation temperatures under various sample compositions. Tau187-RNA CCs were prepared with a fixed [tau]:[RNA] ratio corresponding to the condition of net charge balance. Therefore, the composition of tau187-RNA CC can be determined by [tau] and [NaCl]. Samples were heated at 1 °C/min between T = 15-25 °C, while the turbidity was monitored. The turbidity-temperature data of the heating curves were then fit to a sigmoidal function, so that the cloud point temperature, T_cp_, could be extracted as shown in Fig 2A (T_cp_ was determined from heating curves out of practical utility; T_cp_ from cooling curves is possibly closer to thermodynamic transitions). The experimental cloud-point temperature T_cp_ for CC formation as a function of [tau] and [NaCl] are shown (as points) in Fig 2B and Fig 2C. The experimental data points show that increasing [tau] lowers T_cp_, favoring CC formation, while increasing [NaCl] raises T_cp_, disfavoring CC formation. Such trends were observed at two [NaCl] and two [tau] values, respectively (Fig 2B, 2C). Experimentally, T_cp_ was determined for a range of [tau] and [NaCl] conditions (see Fig S4).

**FIG. 2.**
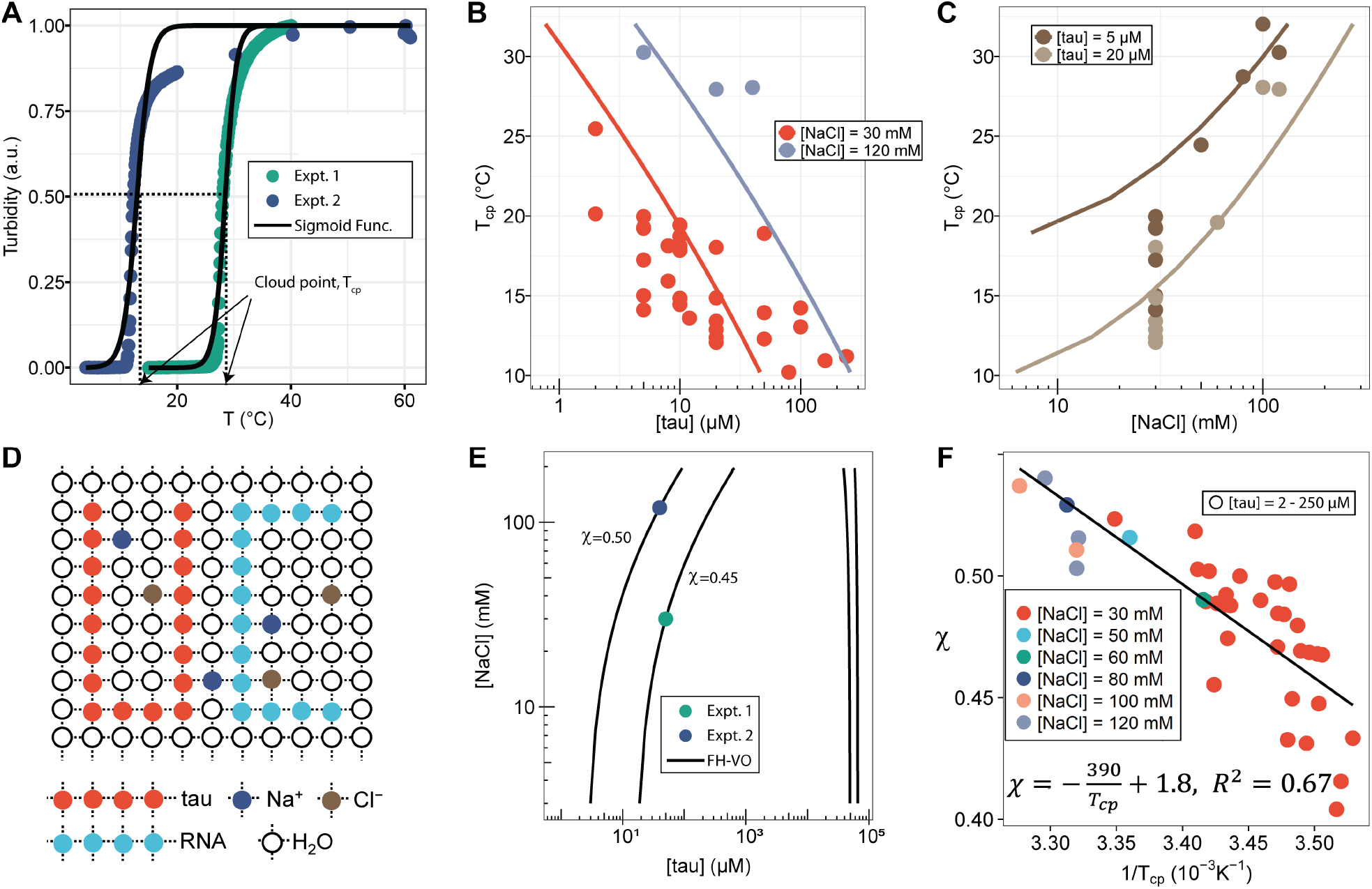
FH-VO modeling of tau-RNA CC. **A.** Turbidity of tau187-RNA CC upon heating (Expt.1 ([tau], [NaCl]) = (50 µM, 120 mM), green dots; Expt.2 ([tau], [NaCl]) = (40 µM, 30 mM), purple dots). Absorbance at λ = 500 nm were normalized and used as turbidity value. Turbidity-temperature data of the heating curves were fitted with a sigmoidal function (solid line) as described in (SI Method), and the temperature at which normalized turbidity reaches 0.5 was assigned to cloud point, T_cp_. **B-C.** Experimental phase diagram (points) showing [tau] vs. T_cp_ and [NaCl] vs. T_cp_ along with the binodal curve generated from fitting the data to the FH-VO model with χ = χ(T_cp_) (solid line) **D.** Diagram of Flory Huggins lattice. Tau and RNA are represented by consecutively occupied lattice sites. **E.** Each experimental condition in (**A**) was independently fit to the FH-VO model (solid lines) to obtain an empirical *χ* value. **E** shows two representative curves. These empirically determined values of χ are shown as points in **F.** The solid line in **F** is a linear regression, generating χ = χ(T_cp_), which is then used to generate the binodal lines in **B** and **C**.

The features of the Tau-RNA CC phase diagram were also investigated by comparing tau187 and tau114. Tau187-RNA CC and tau114-RNA CC were prepared with 20 µM tau187 and 28 µM tau114, so that the total concentration of polymer, i.e. tau and RNA, reaches 0.5 mg/mL. Turbidity was recorded at varying [NaCl]. Similar to the observation with tau187-RNA CC, tau114-RNA CC showed decreasing turbidity at increasing [NaCl] (Fig S5). The [NaCl] values where turbidity reaches 0 were estimated as 131 mM and 150 mM for tau187 and tau114, respectively, implying CC formation is more favorable with tau114 that hence can sustain higher [NaCl]. Based on this, 20 µM of tau187, 131 mM of NaCl and room temperature, 20 °C, were used as the phase separation conditions ([tau], [NaCl] and T_cp_) for tau187, and 28 µM, 150 mM and 20 °C for tau114. These two experimental conditions were used in the next section for comparing the two constructs of tau.

### Flory-Huggins-Voorn-Overbeek Fit to Experimental Phase Diagram

We next used the FH-VO model to fit the experimental data for the tau187-RNA CC system, as is commonly done in LLPS studies. Despite its theoretical deficiencies the FH-VO model is commonly used for its simplicity and ease of implementation. Our system consists of five species: tau187, RNA, monovalent cation (Na^+^), anion (Cl^-^) and water. For simplicity, we explicitly consider only the effect of excess salt, and do not include polymer counterions. The FH-VO model maps these five species onto a three-dimensional lattice (Fig 2D). Each polymer is treated as a uniform chain with degree of polymerization N and average charge per monomer σ. N was taken as the average chain length of the species (1 for monovalent ions). The charge density σ of RNA, monovalent ions and water were set to 1, 1 and 0 respectively. The values for σ of tau187 or tau114 were calculated from the net charge at neutral pH divided by the chain length. The composition of the species is expressed in terms of the volume fraction ϕ of the occupied lattice sites, which are proportional to the molar concentrations (see SI methods for details). As in experiments, tau187-RNA CCs were prepared at fixed [tau]:[RNA] and [Na^+^]:[Cl^−^] ratios. Under these two constraints, the volume fraction of all five species in tau187-RNA CC listed above can be determined with two variables, [tau] and [NaCl], which are experimentally measurable.

Given N, σ, [tau], [NaCl] and T_cp_, the task is to find ϕ_tau_^I^ and ϕ_tau_^II^, the volume fractions of tau in the dilute and dense, coacervate, phase at equilibrium, i.e. the binodal coexistence points. The model and procedure is described in detail in the SI methods. For each experimental observation of T_cp_ determined for a given [tau] and [NaCl] (Fig S4), the FH-VO expression has one unknown parameter, the Flory-Huggins χ term. The Flory-Huggins χ parameter is introduced as an energetic cost to having an adjacent lattice site to a polymer segment occupied by a solvent molecule [61]. Here we take χ to be an adjustable parameter, such that given a suitable expression for χ, the complete binodal curve can be modeled with the FH-VO theory. Consequently, we first solved for χ at each given experimental condition, so that the theoretical binodal curve intersects the experimental data point. Fig 2E shows two representative examples of a theoretical binodal curve (solid line) intersecting a single experimental data point at the given [NaCl] and [tau]. This procedure gives an empirical χ parameter for each experimental data point, as collated in Fig 2F as a function of 1/T_cp_. We then performed to this set of experimental data a least-squares fit of the empirical χ parameter to the form A + B/T (Fig 2F), yielding an expression of the temperature dependence of χ of

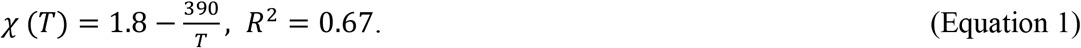

Finally, from this expression for *χ*(T), we computed the binodal curves that establishes the phase coexistence as a function of T_cp_, [tau] and [NaCl], shown as solid lines, only for the dilute phase coexistence for T_cp_ vs [tau] (Fig 2B) and T_cp_ vs [NaCl] (Fig 2C). For the full phase diagram showing both dilute and dense binodal curves see Fig. S7. The experimental data (shown as points) and computed binodal curves both exhibited a decreasing T_cp_ with increasing [tau] and an increasing T_cp_ with increasing [NaCl]. This simply establishes that tau-RNA CC favors higher tau concentrations in the 1-240 μM range and lower ionic strength in the 30-120 mM range tested here. Binodal curves for tau114-RNA CC were also computed, and are compared with tau187-RNA CC, along with experimental data (Fig. S5). Comparison of the two constructs shows that tau114-RNA CC has a lower T_cp_ than tau187-RNA CC, suggesting it is more favorable to phase separation. This qualitatively agrees with experimental observations. Note that the experimental conditions are for [tau] and [RNA] set at charge matching conditions for maximal CC, and thus [tau] and [RNA] are locked relative to each other. When [RNA] falls either well below or well above charge matching condition relative to [tau], it is expected that the LLPS envelope will collapse.

### Field theoretic simulations of a coarse-grained model of tau-RNA complex coacervation

Although the FH-VO model can be brought into agreement with experiment through a judicious choice of χ, it is fundamentally unsound from a theoretical perspective, noticeably because it neglects connectivity between charges on the same chain. This is a severe limitation because it is expected that subtle difference in primary amino acid sequences may have a profound effect on the phase diagram. A particularly appealing alternative to gain insights into the thermodynamics of LLPS is to perform field theoretic simulations (FTS) on a physically motivated polyelectrolyte model (Fig 3), in which each amino acid is represented by a single monomeric unit of length *b* in a coarse-grained bead-spring polymer model. The charge of each segment is unambiguously assigned from the particular amino acid charge at pH 7.0. In addition to harmonic bonds between nearest neighbors, which enforces chain connectivity, all segment pairs interact via two types of non-bonded potentials: a short-ranged excluded volume repulsion and a long-range electrostatic interaction between charged monomers (see Fig 3). We take the polymers to be in a slightly good solvent, meaning that favorable interactions between monomers and solvent cause chain swelling. In such cases, the excluded volume interaction is modeled as a repulsive Gaussian function between all monomer pairs with a strength that increases with solvent quality [62]. Conversely, as solvent quality decreases, the excluded volume repulsion decreases, approaching zero at the so-called theta condition. In the present case we limit ourselves to the case where the excluded volume is positive and small, i.e. a good solvent near the theta condition. Simulations are performed using a single excluded volume strength, *v*, identical for all monomers, which is an input parameter in the model and can be adjusted to parameterize the favorable monomer-solvent interactions. Additionally, the long-range electrostatic interactions are described by a Coulomb potential in a screened, uniform, dielectric background. The length scale of the electrostatic interactions is parameterized by the Bjerrum length *l_B_*, which is the distance at which the electrostatic interactions become comparable to the thermal energy *k_B_T* and is defined as

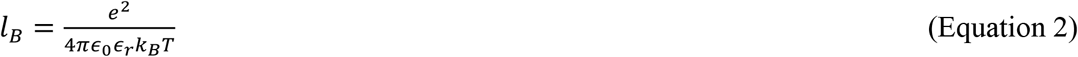

where *e* is the unit of electronic charge, *∈_r_* the dielectric constant (*∈_r_* = 80 for water), and *∈_r_* the vacuum permittivity.

**FIG. 3.**
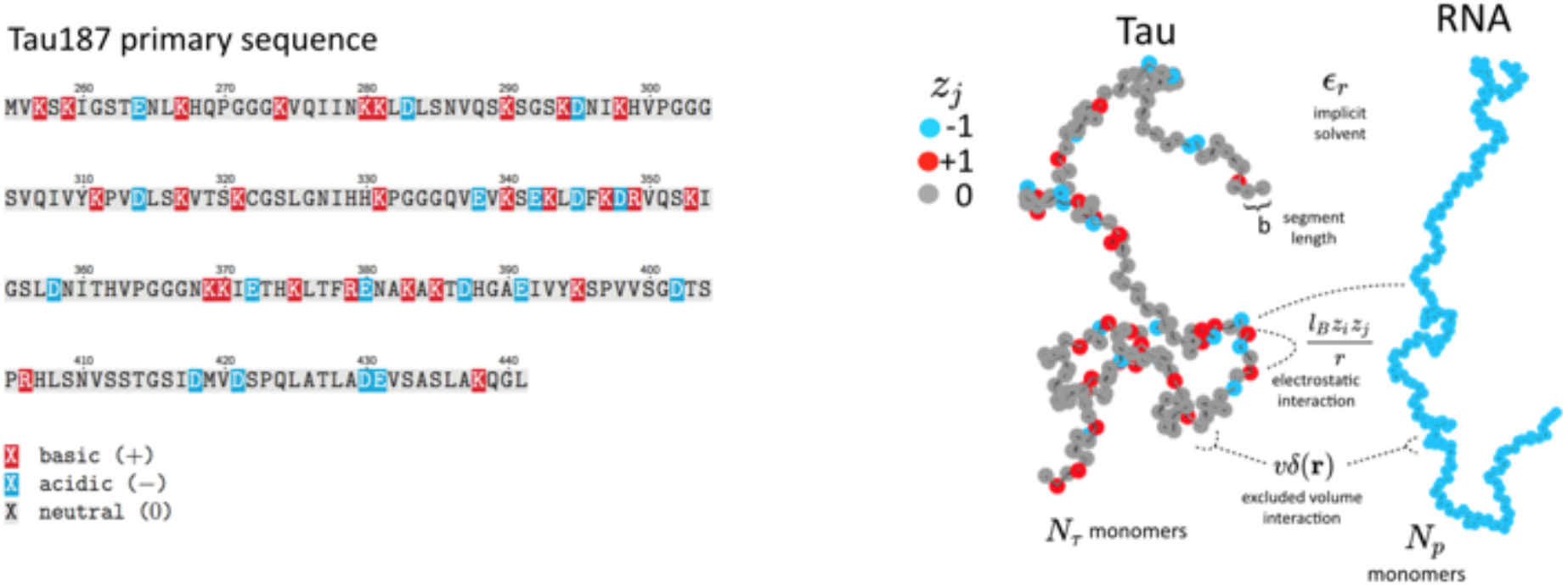
Schematic depiction of the polyelectrolyte model. Tau and RNA molecules are represented as bead-spring polymers with segment length *b* in implicit solvent. Tau is modeled as a polyampholyte with the charge of each monomer determined from the amino acid charge at pH=7. RNA is modeled as a fully charged polyelectrolyte. In addition to chain connectivity, all monomers interact with an excluded volume repulsive potential, and charged monomers interact with a long-ranged Coulomb potential.

The main features of the model used for FTS here are the inclusion of chain connectivity, charge sequence-dependence for the electrostatic interactions based on the primary amino acid sequence of tau, solvation effects which are parameterized by the single excluded volume parameter, *v*, and an electrostatic strength parameterized by the Bjerrum length, *l_B_*. FTS is performed in implicit solvent with a uniform dielectric background. We assume that the polymer chains are in a fully dissociated state, and we do not explicitly represent counter ions. The effect of excess salt is included in our model by introducing point charges explicitly, which engage in Coulomb interactions with all other charged species and repel other ions and polymer segments at short distances by the same Gaussian excluded volume repulsion. By introducing explicit small ions in this manner, we are neglecting strong correlations such as counter ion condensation; however, we are allowing for weak correlations of the Debye-Hückel type. The explicit addition of salt will serve to screen the electrostatic interactions and inhibit the driving force for CC, in agreement with the experiments.

Details of the FTS protocol are described in the SI methods. By performing FTS at various state points and computing equilibrium properties, we first set out to fully explore the parameter space relevant for LLPS in this model. This involves running simulations at different conditions analogous to experiments. For each simulation the thermodynamic state of the system is determined by specifying a particular value for the dimensionless excluded volume parameter *v/b*^3^, the dimensionless Bjerrum length *l*_B_/*b*, and the dimensionless monomer number density *ρb*^3^. Fig 4 shows the final polymer density configuration for two representative simulations at a monomer density of *ρb*^3^ = 0.22 at different thermodynamic conditions (see caption for Fig 4 for details). Although the bulk density is fixed and identical for the two cases, the local polymer density is free to fluctuate. The left simulation box (Fig 4) shows a case where a single phase is favored, indicated by a nearly homogenous polymer density throughout the simulation box (white/blue). This is contrasted by the right simulation box (Fig 4) depicting the case where the system phase separates into a dilute polymer-deplete region (white) and a dense polymer-rich droplet region (red)—the coacervate phase with the color signifying the polymer density. Fig 4 shows that given suitable parameterization, FTS can be used to study complex coacervation of a coarse-grained tau-RNA model. Given this observation, we next map out the full phase diagram in the parameter space of the model while fixing the physical parameters of charge sequence, chain length, and chain volume fractions that are consistent with the experimental conditions.

**FIG. 4.**
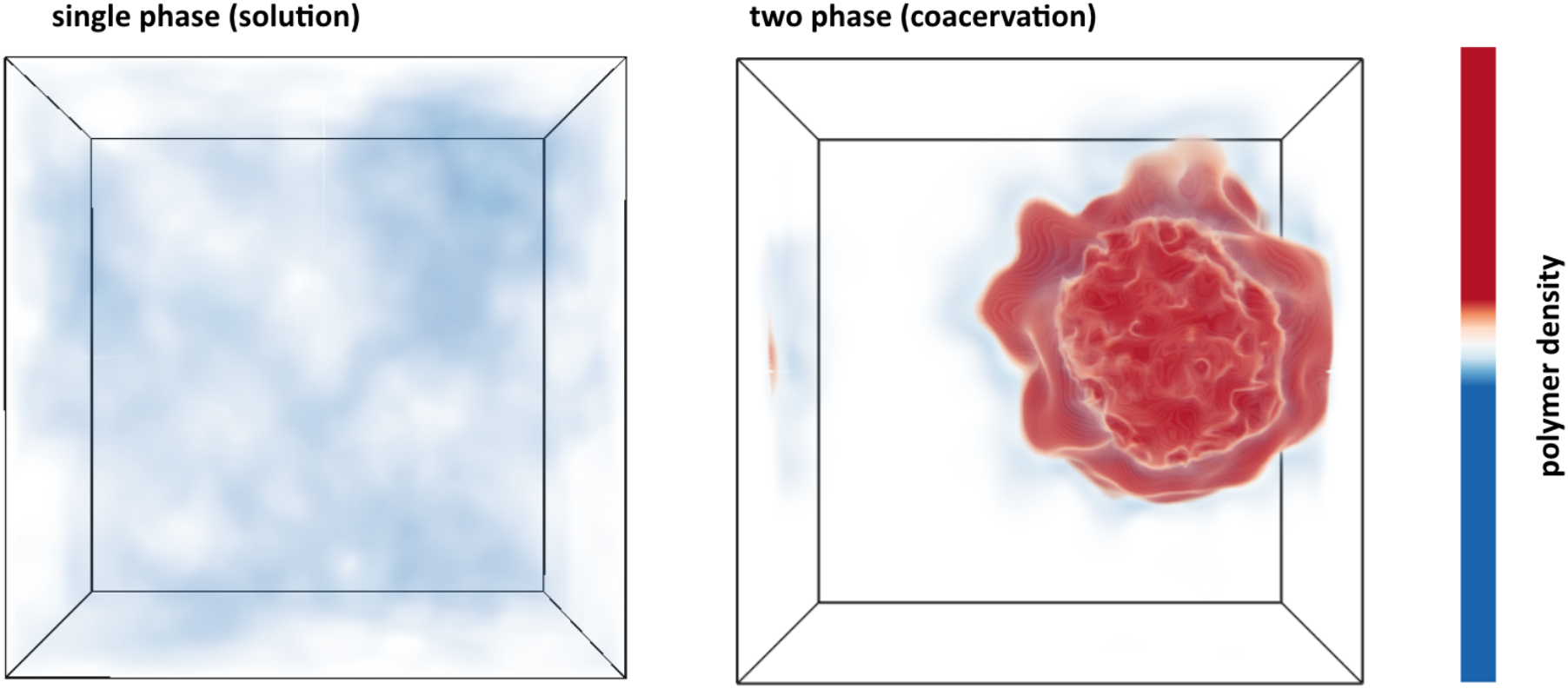
Polymer density field from FTS. Left: Polymer density profile showing a single solution phase for the condition of relatively weak electrostatic strength *l_B_ = 0.16 b* and relatively high excluded volume (good solvent conditions) *v = 0.02 b^3^*. The solution phase is characterized by near homogeneous low polymer density (white/light blue) throughout the entire simulation box. **Right:** Polymer density profile showing complex coacervation upon increasing the electrostatic strength to *l_B_ = 3.25 b* and decreasing the solvent quality by lowering the excluded volume to *v = 0.0068 b^3^*. The two phase region is characterized by a distinct region of high polymer density (dark red) and a surrounding region of low polymer density (white) within the same simulation box. The total polymer concentration is the same in both simulations.

### Field theoretic simulations predict phase equilibria around physiological conditions

The parameters to be explored in connection with phase behavior are the strength of the interactions in the polyelectrolyte model: the excluded volume strength *v* and the Bjerrum length *l_B_*. A direct comparison between FTS and the experimental phase diagram will be deferred until the following section. The phase coexistence points (binodal conditions) for a given value of the excluded volume *v* and Bjerrum length *l_B_* can be obtained by running many simulations over a range of concentrations, and finding the concentration values at which the chemical potential and the osmotic pressure are equal in both phases (see Fig. S10). The procedure is described in the SI, and is repeated for many different *v* and *l_B_* combinations. The resulting phase diagram will be a three-dimensional surface which is a function of *ρ*,*v*, and *l*_B_. In Fig 5A, we show a slice of this surface along the *l_B_* − *ρ* plane with a fixed value of *v* = 0.0068 *b*^3^, and in Fig 5B we show a slice along the *v* − *ρ* plane with a fixed *l_B_* = 1.79 *b* (at *T* = 293 K, Eq. 3). It should be noted that Fig 5 presents the first approximation-free phase diagrams presented in the literature of a theoretical model describing a biological complex coacervate system. From Fig 5, one can see that *l_B_* and *v* have counteracting effects, namely increasing *v* that is caused by increased solvent quality destabilizes the coacervate phase and favors the single phase, whereas increasing *l_B_* that is caused by reduced electrostatic screening favors coacervation, and destabilizes the single phase. The physical interpretation of the trends in Fig 5 is that the actual binodal for the experimental system will depend on two competing features: the solvent quality proportional to *v*, which inhibits coacervation, and the electrostatic strength of the media proportional to *l*_B_ which promotes coacervation.

**FIG. 5.**
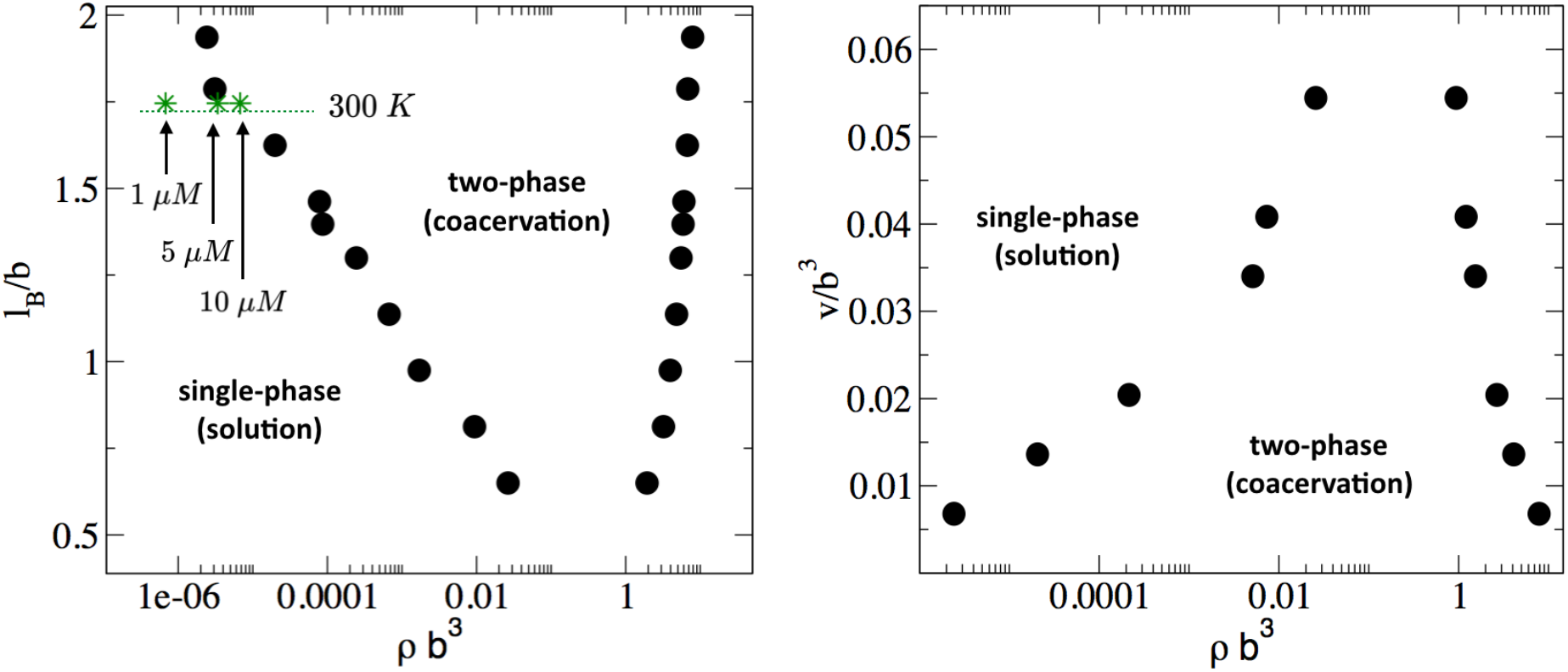
Phase diagram of the Tau/RNA model obtained from FTS. **(Left)** Binodal points as a function of the Bjerrum length at fixed excluded volume of v = 0.0068 b^3^. For comparison three concentrations at 300 K are indicated (arrows) assuming *∈_r_* = 80 and b = 4 Å **(Right)** Binodal points as a function of the excluded volume at fixed Bjerrum length l_B_ = 1.79 b

The FTS- derived phase diagram shown in Fig 5 provides a guide how to experimentally tune the window for complex coacervation by changing the relative contribution of the solvent quality or the dielectric strength. Experimentally, the solvent quality can be decreased by adding crowding agents or by changing the hydrophilic/hydrophobic amino acid composition, while the electrostatic strength can be controlled by the salt concentration. Increasing salt concentration tends to decrease the bare electrostatic strength by screening the charges, and this is predicted to surface along the stabilize the single phase solution mixture against coacervation, in agreement with experimental observation. We explore these ideas further below in the context of tau coacervation *in vivo*.

Despite the simplicity of the coarse-grained description, the model predicts that these two competing parameters, excluded volume vs. electrostatic interactions, are nearly balanced around physiological salt concentration, temperature, and protein concentration. Assuming that the relative dielectric constant for water is *∈_r_* = 80, and that the segment size *b* is approximately equivalent to the distance between *c_α_* carbons, i.e. b ~4 Å, it follows that *l*_B_ = 1.75*b* at 300 K.(*l*_B_ = 0.7 *nm* at 300 K). In the *l*_B_ − ρ plane (shown in Figure 5A), at the cross section of *l*_B_ = 1.75, three points for *ρb*^3^ are indicated that correspond to 1, 5, and 10 *μM* for tau concentrations at 300 K. Here we have implicitly assumed that at physiological temperature and in a crowded cellular environment tau is near the theta condition, and thus *v* is small. This analysis suggests that small modulation in the experimental conditions, such as changes in the temperature or salt concentration, local pH or crowding effects (via the excluded volume parameter v) can readily and reversibly induce complex coacervation *in vivo* under physiological conditions.

### Comparison between simulation and experiment

In the preceding section we presented the phase diagram from FTS explicitly in terms of the model parameters of the excluded volume *v* and Bjerrum length *l*_B_. We now seek to compare our simulation results directly with the experimental phase diagram. This requires knowing precisely how the model parameters depend on temperature. We again take the monomer size *b* to be approximately the distance between the *C_α_* carbons *b* ~4 Å, and assume a dielectric constant of *∈_w_* = 80 for pure water. The relative dielectric *∈_r_* at a given salt concentration is then estimated from Equation 2. With these values, the Bjerrum length *l*_B_ can be estimated from Equation 3 at the experimental cloud point temperature (Fig 6A), which leaves only one unknown parameter *v*.

**FIG. 6.**
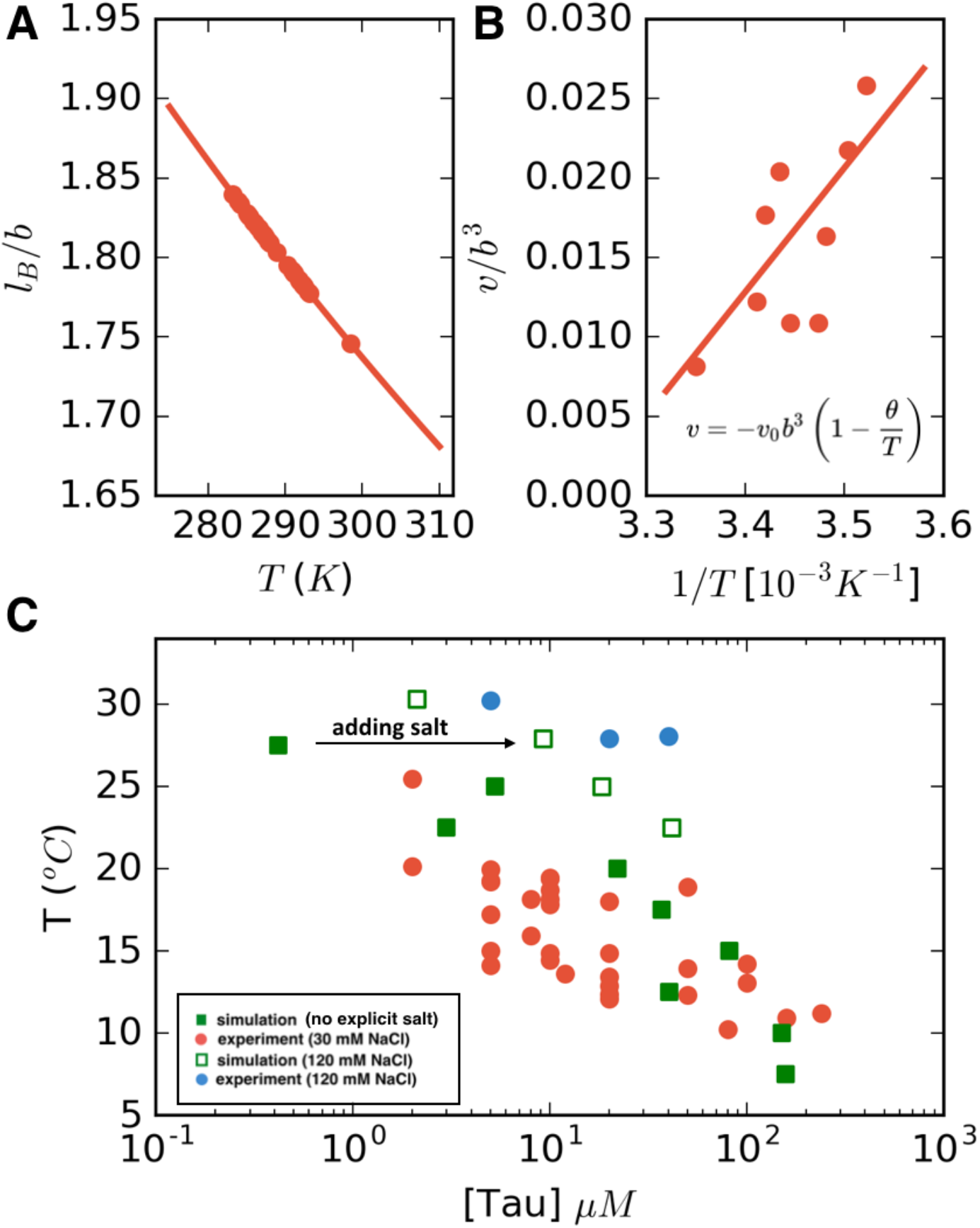
LCST phase behavior from FTS: **(A)** Temperature dependence of the reduced Bjerrum length (ε=80 for water) shown in red. **(B)** Temperature dependence of the excluded volume obtained by adjusting the excluded volume parameter until FTS agrees with a subset of the experimental data (points). The solid line shows a linear fit to the data which is used to obtain the temperature dependent excluded volume for subsequent simulations. **(C)** FTS coexistence points (filled green squares) obtained by using a temperature dependent Bjerrum length (Fig 6A) and excluded volume (Fig 6B). Experiments performed at 20 mM NaCl are shown in red for comparison. Upon introducing excess salt ions in FTS with a fixed concentration of [NaCl] = 120 mM, the binodal shifts upwards (open green squares). For comparison experiments performed at 120 mM NaCl are shown in blue.

The excluded volume is typically taken to be proportional to (1 − *θ/T*) where *θ* is the theta temperature, the temperature at which the chain follows ideal chain statistics [62–64]. For LCST behavior it is customary to introduce the form *v* = −*v*_0_(1 − *θ/T*) where *v*_0_ controls the magnitude of the excluded volume interactions [65]. This form of the excluded volume implies that at temperatures lower than the theta temperature, the excluded volume is repulsive (*v* > 0, meaning a good solvent) and for temperatures above the theta point, the excluded volume becomes attractive (*v* < 0, poor solvent conditions). By adjusting the excluded volume in FTS to fit a subset of the experimental data, (shown in Fig. 6B), we then perform a linear fit to obtain a value of *v*_0_ = 0.25*b*^3^ and *θ* = 309 K. Note that in the range of temperatures considered, the excluded volume remains positive. However, for temperatures higher than θ, when the excluded volume becomes negative, the polymer chain will collapse, consistent with the observation in the literature [66], which showed that tau undergoes a thermal compaction at high temperatures due to entropic factors [63]. In such high temperature regimes a more sophisticated treatment is needed; however, all our experimental conditions remain below this threshold. Having mapped the two model parameters *v* and *l_B_* to the experimental temperature, we can compare directly the FTS with the experimental results (Fig 6C). The calculated FTS data points under the condition of low salt concentration are shown as filled green squares in Fig 6C.

Next, explicit salt ions were introduced as point charges to simulate an excess salt concentration of 120 mM. We make the assumption that the salt is equally partitioned in both phases, and thus the concentration of salt is a constant, allowing us to sweep the polymer concentration at fixed salt concentration to find the phase coexistence points. A more detailed FTS study of salt partitioning performed using a Gibbs ensemble method found that under conditions of nearly charge-balanced polymers, as is the case in the system of this study, the salts are nearly equipartitioned and counterion condensation is not a dominant factor [67]. Simulations performed in this manner with explicit salt are shown as open green squares in Fig 6C. The FTS data clearly demonstrate that the effect of added salt is to stabilize the single solution phase, and to raise the binodal closer to physiological temperature 37 °C, in agreement with experiments (filled red and blue circles Fig 6C). The complementary dense branch of the binodal curve is also predicted from FTS and is shown in Fig. S11.

### Application to cell-complex coacervate co-culture

Looking at the experimental and calculated phase diagrams (Fig 2B, 2C), it is seen that under physiological conditions (T_cp_ ~ 37 °C, [NaCl] ~ 100 mM) it is principally feasible for cells to tune the formation of tau-RNA CCs. This has important implications for studying the physiological roles of tau-RNA CCs, and thus we asked if tau-RNA CCs could indeed exist in a biologically relevant media in the presence of living cells. Both the FH-VO theory and FTS predict that the conditions of high protein concentration, low ionic strength, high temperature and high crowding reagents (acting to lower the excluded volume) would independently favor tau-RNA CC formation. Using these tuning parameters as a guide, we designed several experiments to test the ability for tau-RNA CCs to form in a co-culture with H4 neuroglioma cells. We incubated H4 cells with tau187/tau114-RNA under CC conditions at varying temperatures, polymer concentrations and crowding reagent concentrations. At low polymer concentrations (10 µM tau, 30 µg/ml RNA) no LLPS was observed in the cellular media (Fig 7, first column), where increasing the temperature to 37 °C did not apparently influence the solution phase (Fig 7, first column, first and third row). However, when tau and RNA concentrations were increased (100 µM tau, 300 µg/ml RNA) LLPS could be observed (Fig 7, second column). Further, LLPS could also be achieved by adding an additional crowding reagent (here PEG) to low concentration samples of tau and RNA (Fig 7, third column). As predicted, LLPS of tau-RNA CC was modulated by (i) temperature, (ii) tau and RNA concentration and/or (iii) the presence of crowding reagent PEG (Fig 7). Lowering the temperature to 18°C significantly reduced the number and size of fluorescent droplets, demonstrating that tau-RNA LLPS is indeed tunable by temperature, and demonstrate the biological consequence of the LCST behavior (Fig 7, first and third row). These results were consistently found for both tau187 and tau114 systems. The successful application of FTS for tuning and predicting tau-RNA CCs in cellular media is a first step towards understanding the physiological condition under which tau-RNA LLPS, which follows the CC mechanism, can occur. The conditions described for LLPS here suggests that conditions exist *in vivo* under which LLPS by complex coacervation may be achieved.

**FIG. 7.**
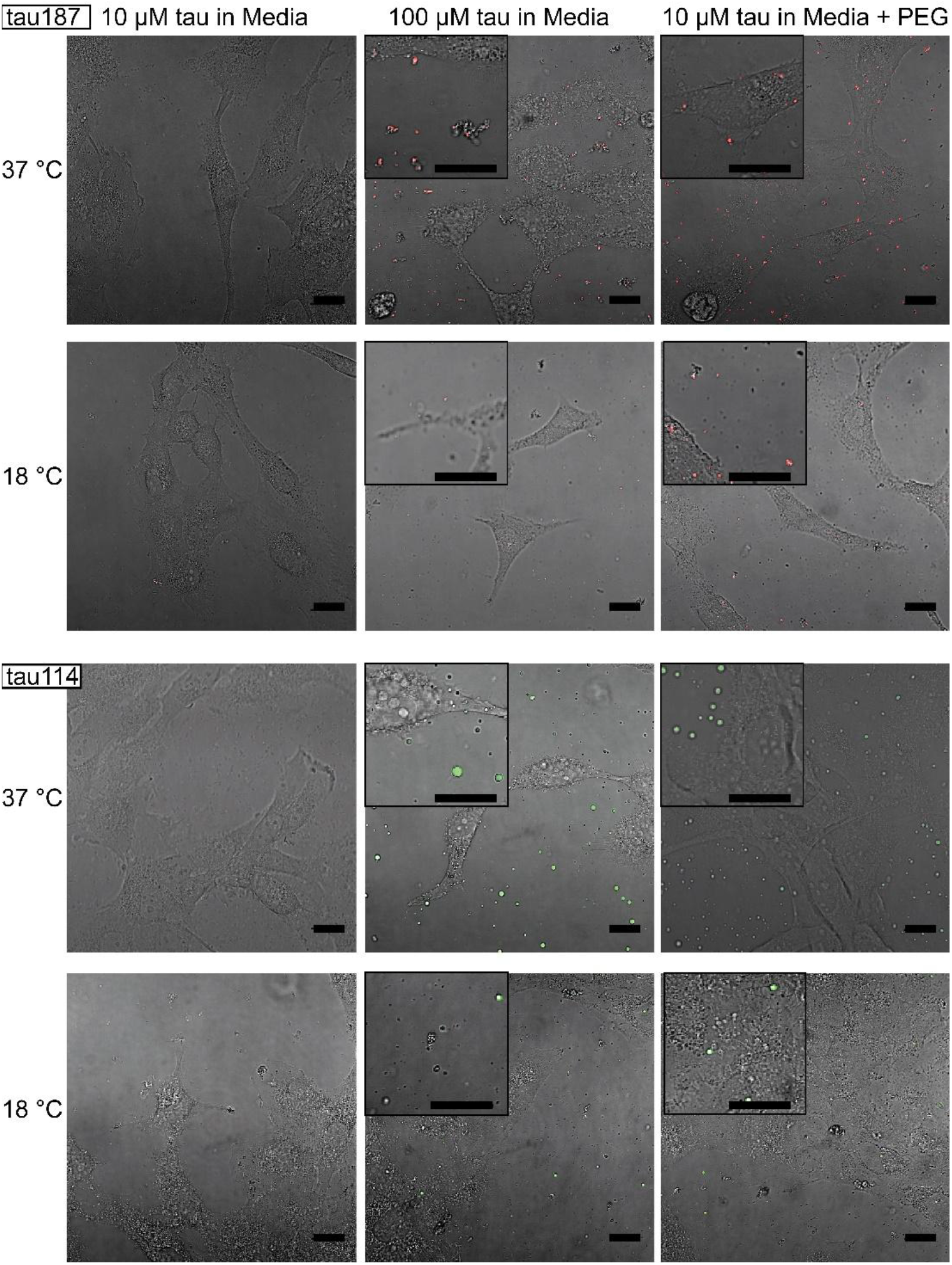
Tuning experimental conditions to catch tau-RNA complex coacervates in presence of living cells. Bright field images and superimposing fluorescence images of tau-RNA CC coculturing with H4 cells, with 10 µM tau (left), 100 µM tau (middle) and 10 µM tau with 10% v.v. PEG (right). Samples at 37 °C (first row) and 18 °C (second row) were images with representative images showing the co-presence of living cells and tau-RNA CCs. Tau187 (Top) and tau114 (Bottom) was used showing tau114 with higher propensity at CC formation. Alexa Fluor 488 was used to prepare fluorescent labeled tau. 3 µg/mL polyU RNA per 1 µM tau was used to prepared samples. Scale bar is 20 µm.

## Discussion

The ability of tau to undergo LLPS via a mechanism of complex coacervation has been recognized in a number of recent publications [21,25] [22]. However, to date, the criteria and physical parameters (specifically, polymer concentration, ionic strength, temperature and crowding reagents) that drive tau-RNA CC has not been rationalized. In this paper, we mapped out the experimental phase diagram for tau-RNA CC, and used theory and simulation to describe the parameter space for LLPS. In what follows, we discuss the relevance of our findings in the context of the physical mechanism of LLPS *in vivo*.

Although the FH-VO model cannot model spatially varying charges along the peptide backbone, we were able to fit the experimental data by treating the Flory-Huggins *χ* parameter as an empirical, temperature-dependent, adjustable parameter. This result highlights the fact that the FH-VO model is adaptable to experimental data. Still, the FH-VO model has limited predictability, and should be seen as a qualitative descriptor of phase separation. In contrast, FTS is an approximation-free analysis that can provide physical insight and predictive information for biopolymers, such as scaling relationships and polymer or protein sequence effects. As shown above, the tau-RNA phase diagram was successfully reproduced by FTS using model parameters that are reasonable estimates of the experimental physical conditions. With reasonable estimates for the parameters in our polymer model (*∈_r_* = 80, *b* = 4 Å), our simulations predict that the lower phase boundary falls in the vicinity of physiological conditions. This finding suggests that FTS can be a powerful theoretical modeling technique to describe and rationalize tau-RNA CC as a competition between short-ranged excluded volume interactions and long-ranged electrostatic interactions.

Consider that we can partition the driving forces of CC as

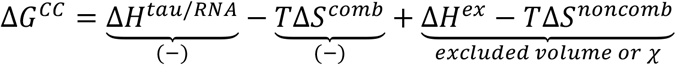

where the first two terms are the negative (favorable) enthalpic contribution from tau/RNA interactions and the ideal entropy of mixing term (which is negative because we are considering CC formation). These first two terms are approximately accounted for in the original VO model, and by themselves predict UCST behavior (see SI). The last two terms introduce an non-ionic excess enthalpic contribution and a nonideal, noncombinatoric entropy that are introduced into the FH-VO model through the Flory-Huggins *χ* parameter, or within FTS through the temperature dependent excluded volume. Given the experimental observation of LCST phase behavior, these terms must be important and we now estimate their value from our model.

Modeling the LCST experimental tau-RNA CC phase diagram using the FH-VO model by invoking an entropic term in the Flory-Huggins χ parameter, or by FTS using a temperature dependent excluded volume, both provide an estimate of the entropic contribution that drives CC formation. The temperature-dependent excluded volume *v* used to describe LCST phase behavior within FTS can be formally related to the Flory-Huggins χ parameter to second order in the *v* = *b*^3^(1 − 2*χ*) [68]. Substituting our empirical excluded volume, we obtain from FTS an interaction parameter χ of the form *χ* = *∈_s_* + *∈_H_*/*T*, with *∈_s_* being a non-combinatoric entropic term and *∈_H_* an enthalpic term. Introducing conventional units (see SI methods for details) gives an unfavorable non-electrostatic enthalpy of phase separation of ΔH^ex^ = 0.23 kJ ⋅ mol^−1^ of monomer, and a favorable noncombinatoric entropy of phase separation of TΔS^noncomb^ = 1.1 kJ ⋅ mol^−1^ of monomer at T = 300 K. For comparison, the empirical χ from fitting the experimental data with the FH-VO model gives ΔH^ex^ = 2.3 kJ ⋅ mol^−1^ of monomer and TΔS^noncomb^ = 3.24 kJ ⋅ mol^−1^ of monomer.

Notably, ΔH^ex^ is small and positive. We hypothesize that the positive, i.e. nonionic, enthalpy value for forming a coacervate phase is due to the requirement of breaking favorable interactions between hydrophilic residues and water that stabilizes the solution phase of tau (ΔH^ex^ = -ΔH^tau/water^). For comparison, the enthalpy of forming a hydrogen bond ΔH_HB_ at room temperature is ~ −8 kJ ⋅ mol^−1^ [69] while the enthalpy of hydration for a polar amino acid ΔH_hyd_ is ~ −60 kJ ⋅ mol^−1^ [70,71]. Given that ΔH^tau/RNA^ for tau–RNA association is negative and tau remains hydrated in the CC state (i.e. tau-water interface is not dehydrated), there has to be a source of penalty in the form of a positive ΔH^ex^ value; the unfavorable ΔH^ex^ associated with tau-RNA CC might come from the loss of hydrogen bonds in the hydration shell from overlapping and sharing of the tau hydration shells in the dense CC phase.

The TΔS^noncomb^ value is also small, positive and of comparable magnitude as ΔH^ex^, making temperature increase a facile modulator favoring tau-RNA CC. Given the positive value of ΔH^ex^ polymer volume fractions for tau-RNC CC, the entropy gain upon phase separation is contributing to the driving force of tau-RNA CC formation (besides the electrostatic correlation energy between the polycationic and polyanionic polymer segments that is the major driving force). Looking to potential origins for positive TΔS^noncomb^, we consider the entropy gain of breaking a hydrogen bond of TΔS_HB_ ~ 6 kJ ⋅ mol^−1^ [69] and the entropy gain associated with the release of a single water molecule from a hydrated surface of ~ 7.5 kJ ⋅ mol^−1^ [72]. Given that our FTS study only considered excess ions, but no counterions, while fully capturing the LCST behavior through the excluded volume, *v*, our results are consistent with the hypothesis that competing hydrophilic/hydrophobic interactions are responsible for the LCST behavior [73–75]. At low temperatures, the attractive interaction between water and hydrophilic residues of the biopolymer stabilize the homogenous phase, but above a critical temperature hydrophobic interactions become dominant, in that it becomes more favorable for water to be released from the polymer surface and hydration shell, and for tau and RNA to associate. In this scenario, the entropy gain comes from the release of bound water into the bulk [70] due to overlapping of the hydration shell of tau upon CC. In the literature, the entropy gain of counter ion release [76–79] or compressibility effects [80,81] have been proposed as origins for the LCST behavior, and as prevalent driving forces for CC [82]. While this study cannot entirely delineate between these possible contributions that are all subsumed into the Flory-Huggins χ parameter or the excluded volume parameter in FTS, we demonstrate that it is not necessary to invoke a specific mechanism, such as counter ion release—the most popular hypothesis, to rationalize LCST driven CC formation. In fact, we performed FTS studies with (and without) explicit excess ions (Figure 6C) observing LCST behavior simply by means of excluded volume and electrostatic considerations and not invoking any counter ion release mechanism to capture the phase diagram of the entropy driven tau-RNA CC. Instead, many factors that globally modulate the excluded volume effects in the biological system of interest and that inevitably modulate the hydration water population, including the hydrophobic effect and crowding, may be considered.

We demonstrated here that tau-RNA CC can be modeled as a coarse-grained polyelectrolyte mixture using equilibrium theory, and revealed the associated driving factors and the different thermodynamic contributions to the phase diagram. However, this finding does not contradict the possibility that tau-RNA complex coacervation is followed by, or even can facilitate, amyloid fibrillization of tau. Comparing our study to previous reports in the literature [21–23,25,83,84], it is clear that tau in fibrils possess dramatically different properties than tau in CCs. In contrast to fibrils, tau-RNA CCs are reversible and tau remains conformationally dynamic – this is because CCs are formed with a stable tau variant, such as the WT derived tau studied here. However, once aggregation-promoting factors are introduced, not only can the thermodynamically stable phase of tau-RNA CC be driven out of equilibrium, but the dense CC phase harboring high tau and RNA concentration may also lower the activation barrier for, and thus facilitate, tau aggregation. Still, tau complex coacervation is a distinct state and fibrilization is a distinct process, where the equilibrium of one does not contradict with its kinetic transformation into the other. Recently, the possibility of the transformation of tau CCs into tau fibrils has been demonstrated [21]. We have independently investigated these questions and find that irreversible transformation can be triggered by doping tau-RNA CC with highly sulfated polysaccharide heparin (Fig. S12). Tau is first driven towards an equilibrium complex coacervate state, from which tau can either re-dissolve into solution state reversibly, or form amyloid fibrils when aggregation driving force is present. However, the mechanism by which the CC state of tau influences the rate of aggregation and/or alters the aggregation propensity of tau is not understood, and will and should be the subject of future studies.

The physiological role of tau-RNA CC as a possible regulatory mechanism or as an intermediate toward fibrilization is an ongoing topic of research. In either case, for tau-RNA CC to be relevant for cellular function LLPS would have to be possible near (certain) physiological conditions. Our *in vitro* experiments found the tau-RNA CC phase diagram boundary to lie near physiological conditions. This suggests that tau-RNA CC can occur *in vivo* upon modulation of parameters, such as the local temperature, electrostatic balance, including local pH, and osmotic pressure. We demonstrate that indeed tau-RNA CC can be achieved in co-culture with living cells. While the coexistence of tau and RNA at low (10 µM) polymer concentrations is not sufficient to drive CC in cellular media, the addition of a molecular crowding reagent is, under physiological conditions (Fig 7). While in this study crowding has been simulated with PEG, many cellular proteins can act as molecular crowding reagents. This data encourages us to speculate that mechanisms that increase the already high concentrations of free proteins and other macromolecular constituents, *not* participating in CC, beyond the normal level within the cell (estimates of 50-200 mg/mL [85]) could be sufficient to promote tau-RNA CC by exerting crowding pressure. Thus, biological mechanisms that increase the concentration of intrinsically disordered and charged proteins and nucleic acids may be potent factors that drive liquid-liquid phase separation in the cellular context. Specifically for the context of this study, high concentrations of tau-RNA are by themselves sufficient to drive CC formation (Fig 7). Given that tau is known to bind and localize to microtubules in the axons of neurons, it is not a stretch to envision a scenario where the local concentration of tau would be highly elevated under certain stress conditions, around regions like the axon initial segment. We proposed at these places in neuron, tau-RNA CCs have a higher probability to be observed. However, even though our calculations and experimental data support a model where tau-RNA CC *in vivo* is possible, whether this actually occurs within the cell depends on many other factors, among them the strength of tau-microtubule binding that compete with tau-RNA CC.

## Conclusion

We report here the first detailed picture of the thermodynamics of tau/RNA complex coacervation. The observation of an LCST phase diagram implies that although electrostatic interactions are key to CC formation, factors that contribute to solvation entropy gain are key to driving liquid-liquid phase separation. We have computed the first approximation-free theoretical phase diagram for tau/RNA complex coacervation from FTS, where we introduced a temperature-dependent excluded volume term. Simulations show a competition between electrostatic strength (parameterized by the salt concentration) and excluded volume (parameterized by the solvent quality). This knowledge can be used to design experiments that perturb this parameter space *in vivo*, as well as predict or understand biological mechanisms that may be favorable towards liquid-liquid phase separation. As a proof of this concept we have shown that by deliberately changing salt concentration, temperature, and solvent quality (by the addition of PEG), we can make tau/RNA LLPS appear or disappear in cellular medium with *live* cells. Interestingly, we find that without any adjustable parameters our simulations predict that tau/RNA is positioned near the binodal phase boundary around physiological conditions. This suggests that small and subtle changes within the cellular environment may be sufficient to induce LLPS in otherwise healthy neurons. Even if the conditions that induce LLPS in the cell is transient, the LLPS state can facilitate irreversible protein aggregation if aggregation-promoting factors are already available, giving credence to the idea that LLPS may play a role in neurodegenerative diseases. However, we speculate that LLPS is reversible in the majority of biological events that drive LLPS, making it hard to observe this state within the cellular context.

## Acknowledgement

We acknowledge insightful and sustained discussions with Dr. Xumei Zhang on LLPS of tau under cellular conditions. JM would like to thank Scott P.O. Danielsen for helpful discussions regarding FTS. The authors acknowledge the National Institutes of Health (NIH) http://www.nih.gov (grant number R01AG05605) received by SH and KSK and the Tau consortium http://www.tauconsortium.org received by KSK and SH. The funder had no role in study design, data collection and analysis, decision to publish, or preparation of the manuscript. JS acknowledges support from the NSF (MCB-1716956). We acknowledge support from the Center for Scientific Computing from the CNSI, MRL: an NSF MRSEC (DMR-1720256). This work was partially supported by the MRSEC Program of the National Science Foundation under Award No. DMR 1720256.

